# pH-Sensitive Amino Lipid-Driven Pore Formation Enables Endosomal Escape of Lipid Nanoparticles

**DOI:** 10.1101/2025.11.15.688617

**Authors:** Akhil Pratap Singh, Kana Shibata, Yusuke Miyazaki, Wataru Shinoda

## Abstract

Lipid nanoparticles (LNPs) have transformed nucleic acid delivery for vaccines and gene therapies, yet their efficiency remains limited by incomplete endosomal escape. While experimental studies have revealed endosomal membrane disruption facilitating cytosolic release, the molecular basis of this process remains poorly understood. Here, we employ molecular simulations to elucidate how LNPs fuse with the endosomal membrane and release their payloads. We identify multiple fusion pathways, with a dominant stalk-pore mechanism. Our simulations reproduce and rationalize key experimental observations, including the transfer of ionizable lipids from LNPs to the membrane, which promotes nucleic acid reorientation and facilitates stalk formation and expansion. Furthermore, lipid shape, pH sensitivity, membrane tension, and nucleic acid encapsulation emerge as critical molecular determinants of endosomal escape efficiency. These findings advance our mechanistic understanding of intracellular delivery and provide a framework for the rational design of LNPs with improved performance for gene therapies and RNA vaccines.

## INTRODUCTION

Nucleic acid-based therapies demonstrate significant promise across various applications, including viral vaccines, protein replacement therapies, cancer immunotherapies, cellular reprogramming, and genome editing.^1–4^ However, the effective intracellular delivery of nucleic acid payloads remains a major challenge due to their inherent susceptibility to enzymatic degradation and poor bioavailability, primarily resulting from their large size and strong negative charge.^3^ Successful gene therapy further requires not only targeted delivery of mRNA to specific cell types but also efficient cytosolic release, where the mRNA can engage the cellular translation machinery.^5^

Among the various delivery platforms developed to address these challenges, lipid nanoparticles (LNPs) have emerged as the most effective non-viral carriers for siRNA and mRNA therapeutics, owing to their high drug-loading capacity, biocompatibility, and tunable properties.^6–11^ The clinical success of LNP-based systems has been proven by the mRNA vaccines developed by BioNTech/Pfizer and Moderna, as well as the siRNA-based drug *Patisiran®*.^12–15^

Despite these advances, key barriers remain to achieving the LNPs’ full potential: ^16^ Although LNPs are readily internalized via endocytosis, they often become trapped within endosomal compartments and are subsequently degraded during endosomal maturation and acidification.^17,18^ Inefficient endosomal escape is widely recognized as one of the principal limitations of LNP-based RNA delivery.^19,20^ For instance, studies have shown that only ~2% of siRNA cargo reaches the cytosol following LNP delivery.^21^ For RNA therapeutics to be effective, endosomal escape must occur prior to degradation. Currently, this process is considered a bottleneck in LNP development.^22^ While viral vectors naturally overcome this challenge by fusing with the cellular membranes through specialized proteins, LNP would offer superior alternatives due to their ability to carry larger payloads, along with improved safety and lower immunogenicity, provided that endosomal escape can be sufficiently enhanced.^23^

Current efforts to improve this process have focused primarily on tuning the lipid composition of LNPs.^11,24,25^ For example, LNP containing cholesterol has been shown to enhance delivery efficiency, likely due to its ability to stabilize high-curvature membrane regions involved in LNP– endosome fusion.^26^ Furthermore, ionizable lipids have proven particularly effective: these lipids remain neutral at physiological pH but become positively charged in the acidic endosomal environment, promoting electrostatic interactions with anionic endosomal membrane (EM) lipids and thereby facilitating membrane destabilization and fusion, leading endosomal escape.^8,27–31^

Despite advances in LNP formulation, relatively few experimental studies using spectroscopic and microscopic techniques have directly investigated the endosomal escape process.^21,32,33^ A recent study revealed that damage to the EM, indicated by galectin recruitment, facilitates the release of nucleic acid into the cytosol.^32^ This study also reported the enrichment of ionizable lipids in EMs and the disintegration of LNPs, both of which contribute to membrane destabilization and disruption, potentially leading to nucleic acid release.

However, these experimental studies have not fully elucidated the precise molecular mechanisms underlying endosomal escape, largely due to inherent challenges of visualizing such dynamic, nanoscale processes at molecular resolution. Molecular dynamics (MD) simulations provide a powerful *in silico* approach for exploring these mechanisms at the molecular level. While all-atom (AA) MD simulations can deliver highly detailed insights, they are computationally prohibitive for systems as large and complex as LNPs. In contrast, coarse-grained (CG) simulations offer a tractable means of probing membrane remodeling, fusion events, and phase transitions at relevant time and length scales.^34,35^

A previous CG study has successfully captured the spontaneous formation of inverse hexagonal phases in DNA-lipoplexes and reproduced experimental features such as DNA spacing and lamellar-to-hexagonal phase transitions upon changes in lipid composition.^35^ This study has even modeled fusion events between DNA-lipoplexes and EMs. However, their broader applicability has been limited by the use of general-purpose force fields such as MARTINI, which, while modular and flexible, lack detailed validation for cross-nonbonded interactions essential for accurately describing such complex systems.

In this work, we address these limitations by conducting CG-MD simulations using the pH-sensitive lipid 1,2-dilinoleyloxy-3-dimethylaminopropane (DLin-MC3-DMA or MC3) containing LNPs. Crucially, we explicitly incorporate pH-dependent protonation states of the lipid to mimic the endosomal environment. Nucleic acids are modeled as 12-base-pair double-stranded DNA (dsDNA; PDB: 1BNA), while we recognize that small interfering RNA (siRNA) differs slightly from dsDNA in nonbonded interactions and strand flexibility, which would be only relevant in the case of longer nucleic acid strands. Our CG model (SPICA force field) was developed with parameterizations derived from AA simulations, enabling us to capture specific nonbonded interactions between LNP constituents and realistic EM components, including phosphatidylcholines (PC), cholesterol (CHOL), phosphatidylethanolamine (PE), phosphatidylserine (PS), and phosphatidylinositol (PI).

Our simulations reveal that protonated MC3 interacts strongly with anionic lipids on the luminal side of the endosomal membrane under acidic conditions, leading to localized membrane remodeling and the formation of non-bilayer structures such as stalk intermediates. These structural changes drive fusion-pore formation and subsequent membrane rupture, facilitating the release of nucleic acid payloads into the cytosol. Furthermore, our findings reveal that amino lipids play a crucial role in enabling stalk expansion, which ultimately leads to pore formation and the release of dsDNA. Additionally, our results emphasize the importance of lateral membrane tension in EMs, as well as the spatial distribution of nucleic acids within LNPs, in promoting efficient release of the payload.

Overall, this study provides molecular-level insights into the endosomal escape mechanisms of LNPs, offering rational design principles for developing next-generation delivery systems with enhanced therapeutic efficacy.

## RESULTS

### Lipid-DNA Interactions

To enable LNPs simulations with a model EM, we first determined the nonbonded interaction parameters between nucleic acids and EM lipids that were not available in the SPICA molecular library. Notably, the SPICA force field (FF) already contains all the inter- and intramolecular interaction parameters for MC3 lipids, CHOL, PC, and nucleic acid (dsDNA) (Fig. S1A).^34,36–38^ However, the nonbonded interactions between MC3 or dsDNA and EM lipids (Fig. S1B)—PS and PI—were missing in the SPICA FF. In this study, we explicitly optimized these parameters to reproduce reference data from AA simulations. These data included the surface tension of binary lipid monolayers, density profiles of CG sites, 2D radial distribution functions (RDFs) between the centers of mass of the lipids in the binary monolayers, and the free energy profiles of nucleobases and deoxyribose (DERB) across PS and PI membranes. The optimized CG parameters closely reproduced the AA reference data (Figs. S2, S3, and Table S1).

### MC3 lipid programmed LNP-Endosome Fusion Pore Formation and Expansion

To investigate the molecular origin of endosomal escape mechanism, including the puzzling EM damage observed experimentally,^32^ we performed unbiased CG MD simulations of hydrated EM with solvated LNPs (Fig. 1A), using pre-equilibrated structures described in the Methods section. We simulated four replicas, each for 20 to 30 μs, with slightly different ion concentrations and varying ratios of charged to neutral MC3 lipids in the LNPs. All simulated replicas and their compositions are summarized in Table S2. Replicas 1 and 2 underwent successful fusion of the LNPs with the EM, resulting in payload delivery into the cytosol. Notably, replica 2, at physiological ion concentrations (150 mM NaCl) and containing LNPs with a mixture of charged and neutral MC3 lipids (representing slightly higher pH~5 conditions), yielded qualitatively similar results to those described here. In contrast, replica 3 and replica 4 exhibited only membrane fusion activity but did not show transfection within the 30 μs simulation time. We will discuss this further below.

**Figure 1.**
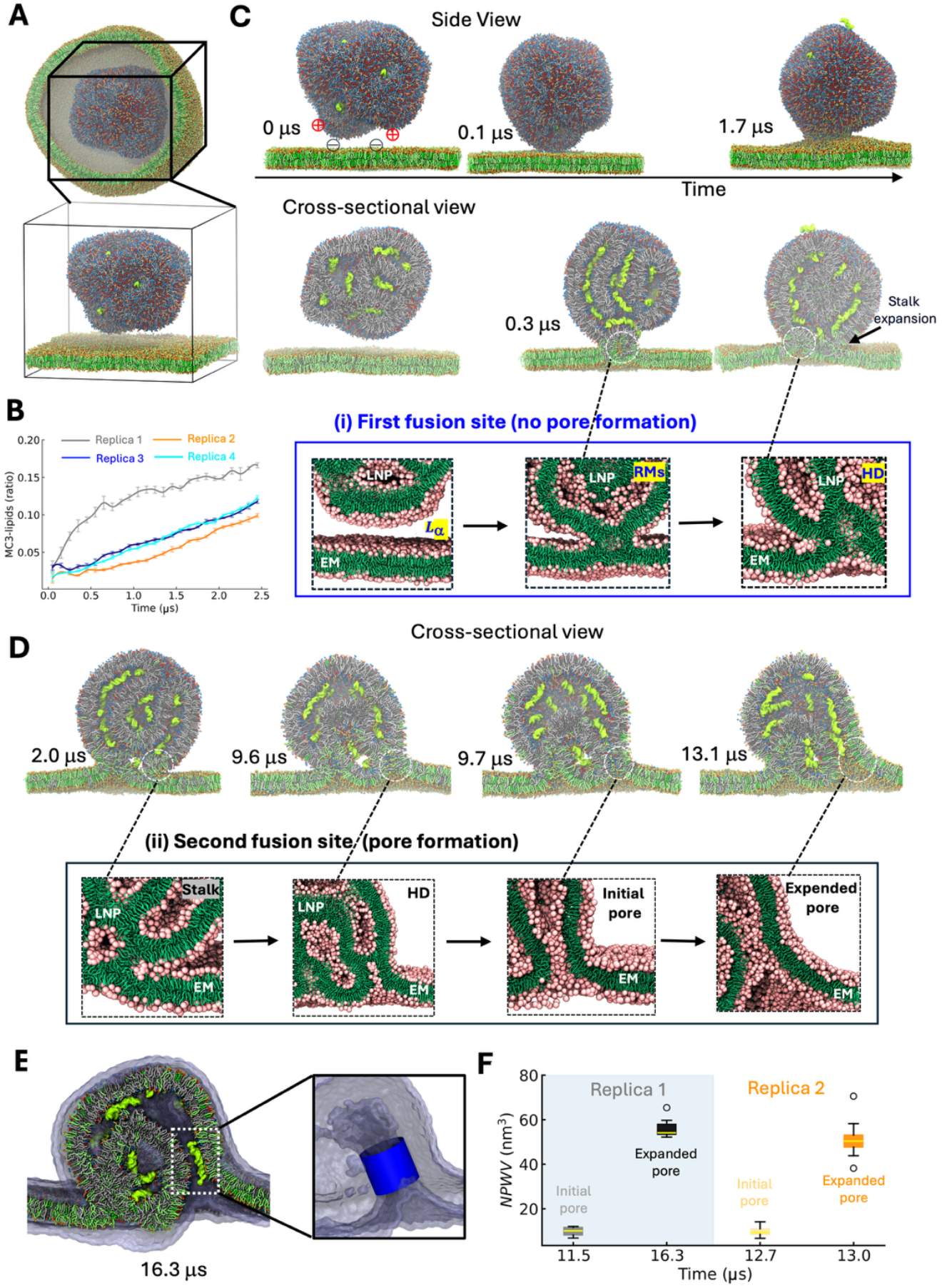
LNP-endosome fusion pore formation and expansion. *(A)* Schematic representation of an LNP entrapped within an endosome. Inset: CG representation of the LNP-EM system. Color scheme: Different lipid types are shown in distinct colors. Lipid headgroups, tails, and dsDNA molecules are depicted using van der Waals (VDW), Licorice, and QuickSurf (iso-surface) representations, respectively. For clarity, water and ions are omitted. (*B*) Time evolution of the fraction of migrated MC3lipids in the EM after stalk formation, showing rapid translocation in all replicas. The fraction of migrated MC3 lipids was calculated relative to the total number of MC3 molecules. Error bars represent standard errors estimated by block averaging. *(C, D)* Representative snapshots showing the time evolution of the LNP-EM system during MD simulations at *(C)* first fusion site and *(D)* second fusion site, leading pore formation and expansion. The color scheme is the same as in panel *(A)*. Inset: Representative snapshots of membrane fusion stages: *(i)* At the first fusion site, two closely apposed bilayers form an HD via an intermediate reverse micellar (RM) phase. *(ii)* At the second fusion site, radial expansion of the stalk and subsequent HD formation initiate and enlarge a small fusion pore. Color scheme: Pink for CG atoms of lipid headgroups, green for lipid tails. For clarity, water, ions, and dsDNA molecules are omitted. (*E)* Cross-sectional view of the LNP-EM complex at 16.3 μs, showing the solvent-accessible surface area (SASA) represented by iceblue color. An inset highlighting a cylindrical region at the narrowest pore for counting water molecules. (*F*) The semi-quantitative estimation of the narrowest pore water volume (*NPWV*) reveals significant pore expansion occurring in both Replica 1 and Replica 2.

Analysis of MD trajectories revealed that LNPs approached the EM within tens of nanoseconds (Fig. 1C). This is consistent with the expectation that strong electrostatic interactions between the cationic MC3 lipids and anionic PS/PI lipids drive the initial contact. However, fusion did not proceed at this stage due to insufficient interfacial dehydration, a known kinetic barrier to membrane fusion.^39^ To address this, we modified the initial setup to include a dehydrated interface, with one corner of the LNP oriented toward the EM, resulting in successful stalk formation (Fig. 1C).

Interestingly, LNP-EM fusion followed two distinct pathways, both culminating in a hemifusion diaphragm (HD) state,^40^ in which the outer leaflet of the LNP and EM merged and exchanged lipids (Figs. 1C, D, S4). At the initial fusion site, a lipid aggregation such as a reverse micelle (RM) formed and subsequently evolved into the HD structure (inset of Fig. 1C). No further changes occurred at this site. However, it is worth mentioning that we observed the splay of MC3 lipids in the EM during HD formation across all replica (Fig. 1B, S5), consistent with experimental report.^32^ This is likely due to the unique structure of MC3 lipids, which feature charged headgroup and polyunsaturated tails, causing greater disruption at the fusion site compared to conventional lipid bilayer or vesicles and typical membrane fusion stalk formation (Fig. S6).

Importantly, the migration of cationic MC3 lipids into the EM creates an ionic imbalance in the LNPs. This imbalance triggers the rearrangement and accumulation of dsDNA near the fusion site, which we will discuss further in the next section. Consequently, this dsDNA reorganization promoted the formation of an expanded stalk-like intermediate, which rapidly progressed into the formation and expansion of a fusion pore (Fig. 1D, E). This fusion pore enabled the release of the dsDNA payload into the cytosol. Notably, we observed distinct time evolutions for pore formation and expansion in both replica 1 and replica 2 (Fig. 1D, S4, 1F).

These findings are consistent with the stalk-pore fusion model widely supported in prior membrane fusion studies.^41–43^ All things considered, protonated MC3-lipid maps out an active fusion site for pore creation, presumably causing EM damage, in addition to the LNP and EM interaction, in agreement with experimental reports.^32,44^

### Cationic-MC3, PE, PS Lipids Facilitate Membrane Fusion

LNP transfection efficiency is significantly influenced by the shape and arrangement of lipid components.^45^ To understand how different lipids affect fusion pore formation, we firstly visually analyzed lipid migration and sorting during fusion (Fig. 2A). We then quantified the number of lipids transferred between the LNP and EM. Lipid transfer from the LNP to the EM occurred in the order: MC3 > CHOL > DSPC. Conversely, lipid transfer from the EM to the LNP followed the trend: PI > PS > PC > PE > CHOL (Table S3). These trends were further supported by measurements of lipid orientation relative to the membrane normal. (Fig. S7).

**Figure 2.**
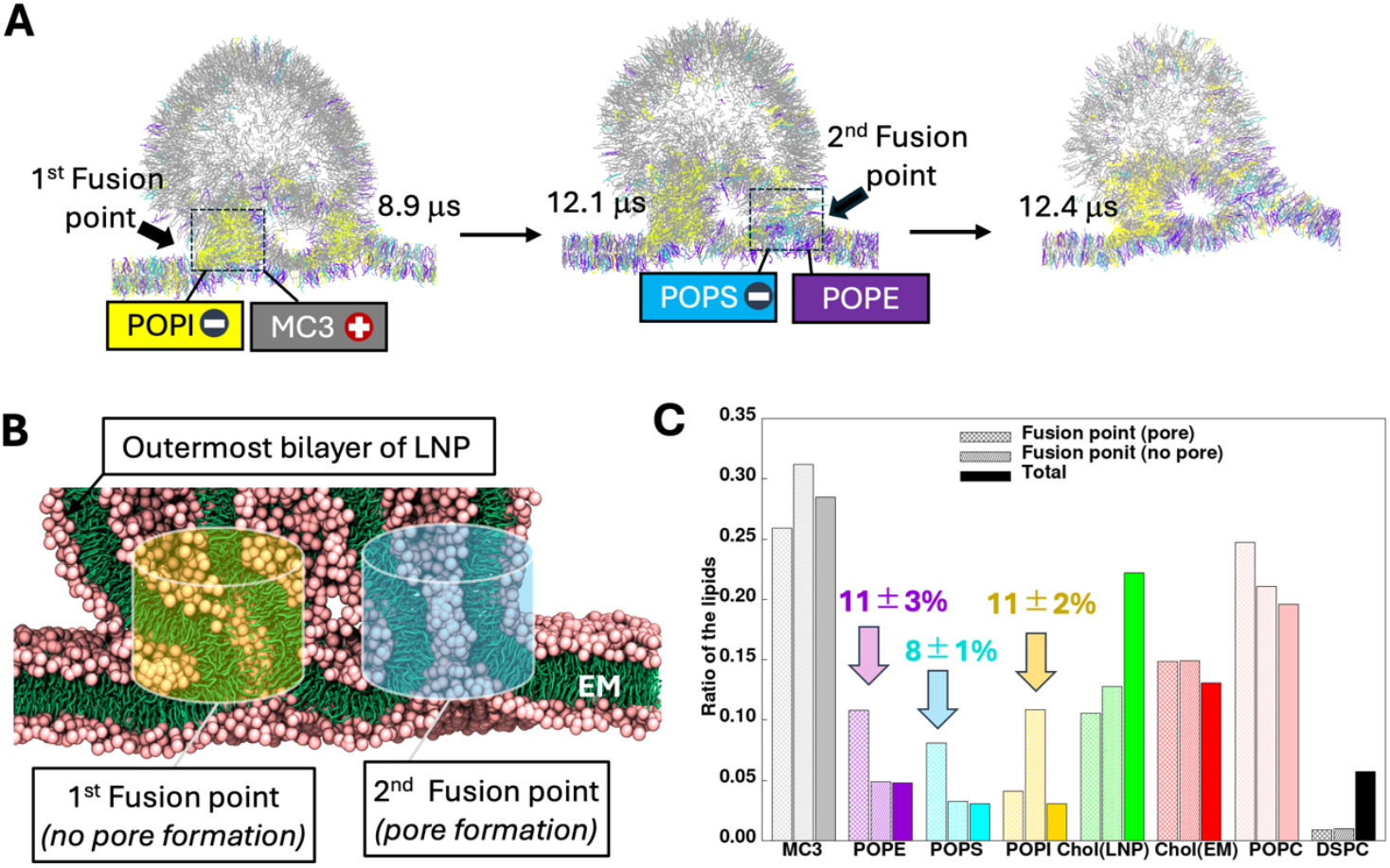
Structural relaxations at fusion points and lipid sorting. *(A)* Representative snapshots showing the time evolution of the MD simulation of LNP-EM, highlighting lipid sorting. Color scheme: gray for MC3, purple for POPE, cyan for POPS, yellow for POPI, and green for cholesterol. For clarity, water, ions, and dsDNA molecules are omitted. *(B)* Cross-sectional view of the fusion points at 12 μs. Lipid compositions at both fusion points are calculated using a cylindrical region, as illustrated. *(C)* Comparison of lipid compositions at the fusion points over the time interval from 11.8 μs to 12.1 μs, based on the cylindrical fusion region shown in panel (B). Errors were estimated using block averaging.

To quantify lipid sorting at the fusion site, we examined the local lipid composition using a cylindrical region centered on the pore (Fig. 2B, C). PE and PS lipids comprised 11% and 8% of lipids, respectively, at the active fusion site, while PI lipids dominated the blocked fusion site. As expected, negatively charged PI and PS lipids were enriched at fusion sites due to electrostatic attraction to the cationic MC3 lipids.

Our results suggest that cone-shaped lipids, such as PE and PS, preferentially accumulate at successful fusion sites, lowering the energy barrier for pore formation, consistent with prior reports.^46–48^

### pH-Sensitive MC3 Lipids and EM Curvature Modulate dsDNA Release

Next, we examined how protonation states of MC3 lipids influence fusion pore dynamics in response to pH changes (Fig 3A). We emphasized the two major scenarios. In the first scenario, all MC3 lipids remained protonated, resulting in slow fusion pore expansion and limited dsDNA release (Fig S8, Movie S1, S2), likely due to strong electrostatic interaction among cationic MC3, and negatively charged dsDNA molecules. In the second scenario, as a consequence of pore formation, we mimicked cytosolic influx into the endosome by deprotonating 64% of MC3 lipids at ~17 µs, consistent with prior experimental estimates.^31^ This charge reduction weakened electrostatic interactions, facilitating pore enlargement and enhanced dsDNA release (Fig 3B, C, Movie S3, S4).

**Figure 3.**
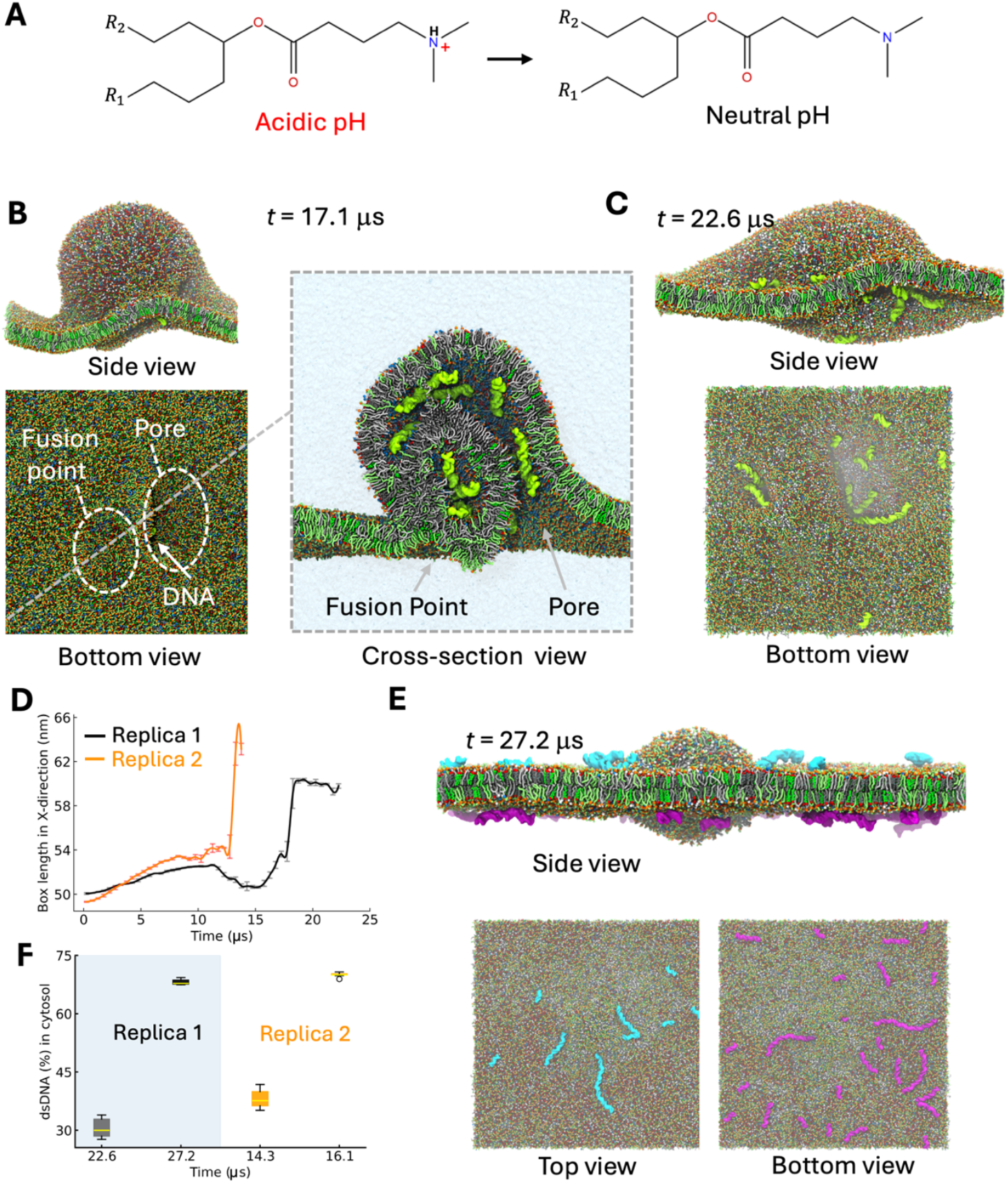
dsDNA release is regulated by pH-sensitive MC3 lipids and lateral tension. *(A)* Molecular structures of the MC3 lipid headgroup at acidic and neutral pH. *(B, C)* Representative snapshots of a single EM-LNP fused complex after randomly replacing 64% of the cationic MC3 lipids with neutral ones at *(B)* 17.1 μs and *(C)* 22.6 μs. The color scheme is identical to that in Fig. 1C. The translucent ice blue represents the water phase. For clarity, ions are omitted. *(D)* The time evolution of the simulation box length in the X-direction for both replicas experiencing pore formation is presented. Error bars on the curves are estimated through block averaging. *(E)* Representative snapshots showing the final configuration of the simulated system after applying lateral tension, which accelerates dsDNA release into the cytosol and leads to closure of the membrane pore at 27 µs. The lipid color scheme is the same as in Fig. 1C. Cyan indicates unreleased dsDNA molecules adsorbed to the inner leaflet of the EM; purple indicates dsDNA molecules released into the cytosol. (*F*) An applied lateral tension to EM at 22.6 μs in Replica 1 and at 14.3 μs in Replica 2 resulted in an approximate 40% increase in dsDNA release in the cytosol, rising from 30% to 70%.

To quantify this effect, we monitored simulation cell size in the lateral direction (Fig 3D), revealing significant EM expansion following deprotonation, consistent with increased lipid migration. These results highlight the critical role of MC3 protonation in regulating pore dynamics and nucleic acid release, in agreement with experimental observations.^11,33^

Despite this, dsDNA release plateaued after 22 µs in replica 1 and 14 µs in replica 2. To further enhance release, we adopted tension-induced fusion as implemented in previous studies.^49,50^ It is important to note that such implementation of tension in EM could also be considered as its intrinsic curvature effect. We thus applied a lateral tension to the EM at 22.6 μs (Fig. 3C) in replica 1 and at 14.3 μs in replica 2, as mentioned in the method section, and continued the simulations. This intervention dramatically improved dsDNA release, with 70% of encapsulated dsDNA entering the cytosol (Fig. 3E, F). These results suggest that membrane curvature and associated tension significantly enhance transfection efficiency.

### Nucleic Acid Distribution Within LNPs Influences Release Efficiency

Finally, we investigated how the spatial distribution of dsDNA within LNPs, resulting from MC3 migration in EM, affects release efficiency. Visual analysis of MD trajectories revealed that prior to fusion, several dsDNA molecules protruded from the LNP surface (Fig. 4A). After fusion, most dsDNA molecules accumulated beneath the LNP’s outermost bilayer (Fig. 4B). Fusion pore formation occurred at regions where the outer bilayer of the LNP contacted the outer leaflet of the EM, leading to cytosolic release of dsDNA (Fig. 4C). In contrast, dsDNA located at the LNP surface failed to escape, as this region fused with the inner leaflet of the EM (Fig. 4E). It is also worth mentioning that the distribution of dsDNA molecules in the LNP-EM fused complexes of replica 3 and 4, which did not undergo pore formation, differed somewhat from the replicas where pore formation occurred (Fig. 4F–I, S9). In the replicas without pore formation (Fig. S10), most dsDNA remained linked to the molecules protruding from the LNP surface. Subsequently, these molecules aligned along the principal axes of the dsDNA and migrated toward the aqueous phase on the luminal side (Fig. S10). However, no DNA was released from the complex into the cytosol during the 30μs time interval, presumably resulting in a more relaxed system.

**Figure 4.**
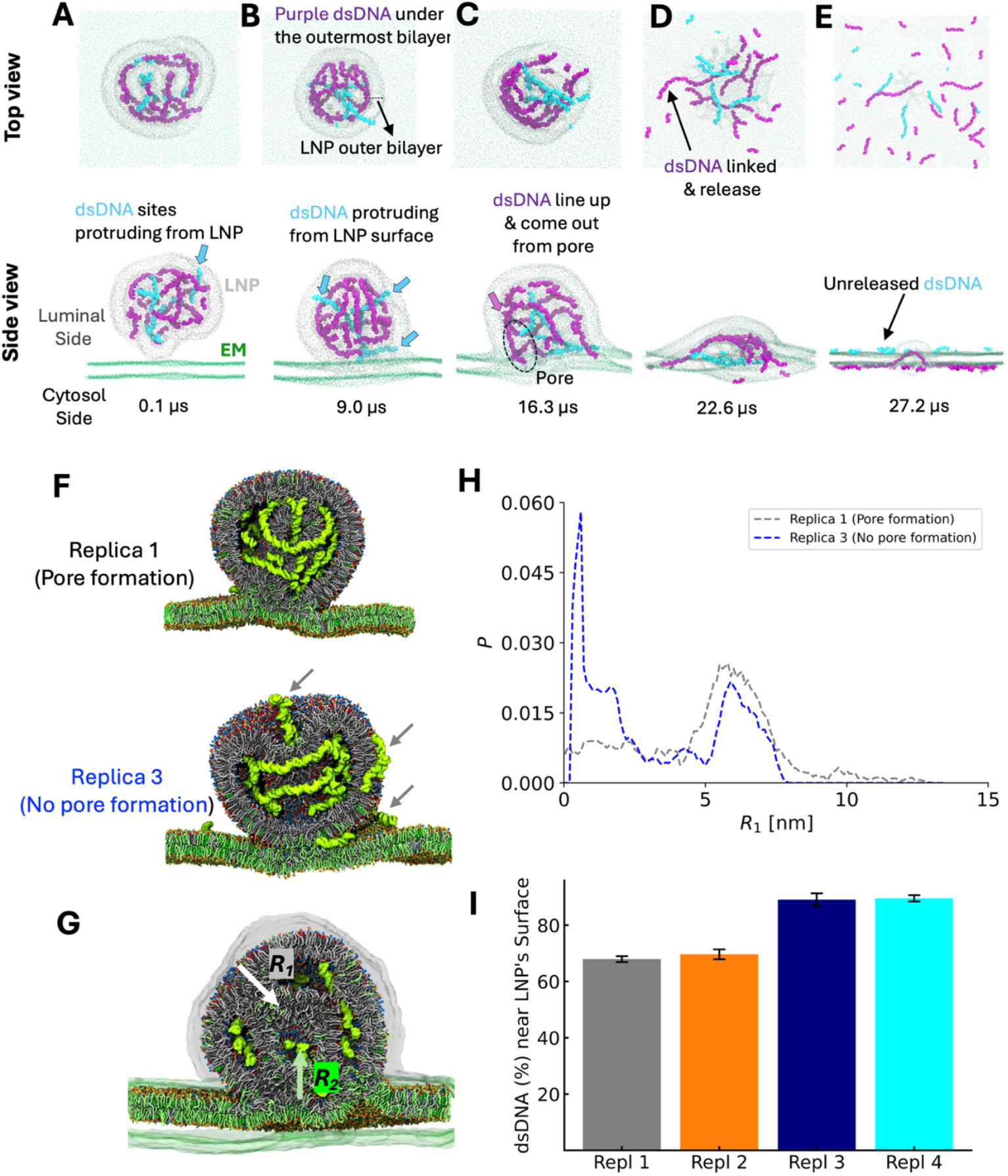
The LNP-EM complex highlights released and unreleased dsDNA molecules, as well as distinct dsDNA distributions in replicas with and without pore formation. *(A, E)* Time evolution of the MD simulation of the LNP-EM complex, highlighting released and unreleased dsDNA molecules in the replicas with pore formation (Replica 1). Gray and green spheres represent the lipid headgroups of the LNP and EM, respectively. The dsDNA color scheme is the same as in Fig. 3E. *(F)* Representative snapshots of a single EM-LNP fused complex in replicas with and without pore formation at ~9 μs. The color scheme is identical to that in Fig. 1C. The arrows in Replica 3 indicate the migration of dsDNA into the aqueous phase on luminal side. For clarify, results from Replica 2 (with pore formation) and Replica 4 (without pore formation) are omitted, as they are similar to those shown here. *(G)* Cross-sectional view of the LNP-EM complex with the solvent-accessible surface area (SASA). Light gray and green transparent regions represent the SASA of the LNP and EM, respectively. *(H)* Probability distributions of dsDNA in the LNP-EM fused complex at ~8 μs from the LNP surface toward its interior. The distance from the LNP surface, *R*_1_, is defined as the inward projection of **R**_1_ = R_i_ − r_SASA_ along the LNP surface normal, shifted by the radius of the SASA probe particle. (*I*) The dsDNA amount within 6 nm (1/4^th^ the diameter of the LNP) of the LNP surface, at around 8 μs, shows an increase by about 20% in replicas 3 and 4, which did not show pore formation, unlike others.

These results indicate that full encapsulation of nucleic acids beneath the LNP bilayer enhances release efficiency, whereas DNA associated with the surface inhibits transfection. Additionally, successful pore formation and transfection require proper alignment of dsDNA molecules towards the fusion site, rather than protruding outward. Overall, our findings suggest that optimizing the internal localization of nucleic acids is crucial for maximizing delivery efficiency.

## DISCUSSION

In this study, we employed CG MD simulations to investigate the endosomal escape mechanism of LNPs and to elucidate the molecular details of their interaction with endosomal membranes, extending beyond the limitations of current experimental methods. While the molecular mechanisms governing nucleic acid transfer from lipoplexes has been extensively studied,^35^ our work distinguishes itself by explicitly modeling the endosomal membrane and a siRNA-LNP therapeutic analog of *Patisiran*^*®*^, using a consistent CG framework rather than the modular “building-block” approach commonly adopted in multiscale modeling. This approach was motivated by two key considerations in addition to exploring EM damage as observed experimentally:^32^ first, the importance of detailed endosomal escape mechanisms for the clinical efficacy of *Patisiran*^*®*^, and second, the potential to leverage these insights for the rational design of more efficient nucleic acid delivery systems.

Our simulations successfully captured LNP-endosomal membrane fusion events, including multiple fusion sites leading to dsDNA transfection. We observed that the fusion pore formation represents a plausible route for endosomal escape, consistent with experimental findings.^32^ However, not all fusion sites developed into functional pores. This suggests a two-step free energy landscape: (1) lipid rearrangement during stalk formation, and (2) transition from HD to pore formation—both consistent with prior studies.^43^ At the initial fusion site, the process stalled at an amorphous HD structure, where multiple membranes from the LNP and endosomal membrane remained entangled, likely due to the high energetic cost associated with PI-rich regions. In contrast, the second fusion event, involving stalk expansion, proceeded through the stalk–HD– pore pathway, facilitating dsDNA release. Notably, our results also revealed that the charged state of MC3 lipids and their polyunsaturated tails disrupted the fusion site, promoting MC3 migration from the fusion site into the EM. This migration drove nucleic acid reorientation within the LNP, leading to dsDNA accumulation at the fusion site, which facilitated stalk expansion and ultimately pore formation.

We further demonstrated that the lipid structure and composition at the fusion site critically influence pore formation and transfection efficiency, supporting previous studies.^47,51^ Cone-shaped lipids such as PS, PE, and cationic MC3 preferentially accumulated at successful fusion sites, lowering the energy barrier for pore formation. In contrast, PI-rich regions, despite facilitating initial fusion due to strong electrostatic interactions, failed to complete the pore formation step.

Next, we examined the effect of pH-sensitive lipid protonation states on fusion dynamics. Initially, under acidic conditions, protonated MC3 lipids promoted LNP-endosomal membrane interactions via electrostatic attraction. Upon pore formation, partial deprotonation of MC3 lipids, mimicking cytosolic pH, reduced these interactions, facilitating rapid pore expansion and nucleic acid release. These observations highlight the significance of dynamic charge regulation in MC3 lipids for regulating endosomal escape efficiency.

Although we could not simulate a fully enclosed LNP within a curved endosome due to computational limitations, we investigated the role of membrane curvature by applying lateral pressure to the endosomal membrane. This intervention, which mimics curvature-induced stress, substantially enhanced dsDNA release. Our findings suggest that membrane curvature and local tension may be critical for achieving complete nucleic acid delivery.

Moreover, our results highlight the significant influence of the spatial distribution of dsDNA molecules within LNP-EM fused complexes, primarily caused by the migration of MC3 into the EM. In our simulations, the two replicas that contained dsDNA molecules beneath the LNP bilayer successfully transfected most of the dsDNA, except for those located at the LNP surface. In contrast, the other two replicas, which did not undergo pore formation, showed rapid leakage of dsDNA during rearrangements within the LNP-EM complex. This leakage prevented the stalk expansion and pore formation. This issue likely stems from the small size of the LNPs, which increases the risk of nucleic acid leakage.^52^ Therefore, LNP size indirectly influences transfection success.

Importantly, these simulation-based insights align with experimental observations from microscopic study^32^ which have revealed transient pore or EM damage and stepwise fusion pathways similar to those observed here. Moreover, LNP disintegration, MC3-lipids migration from LNP to EM, also impact the endosomal escape process, consistent with experimental observations.^32^ Previous simulation studies also support the strong dependence of membrane fusion behavior on local lipid composition and mechanical stresses.^43,47^

Nonetheless, our study has limitations. As with any CG-MD approach, discrepancies exist between simulation and experimental conditions, including differences in RNA type (siRNA vs. mRNA), RNA-to-lipid ratios, salt identity and concentration, and the absence of natural endosomal curvature and proteins. Additionally, the sampling remains limited relative to the complexity of biological membranes. These factors could influence the kinetics and pathways of fusion. Future studies should systematically vary these parameters and incorporate diverse lipid compositions to better understand the energetics and topological constraints governing LNP-endosome fusion.

In conclusion, we present a detailed in silico investigation of LNP fusion with a model endosomal membrane, offering mechanistic insights into the endosomal escape process. Our results pave the way for the rational optimization of LNP formulations. Furthermore, our simulations provide interesting insights beyond the specifics of pore formation, which presumably cause EM damage. It also reproduces and rationalizes experimental observation, which validates a posteriori the quality of our force fields and simulation parameters. Thus, future work could extend these findings by incorporating cell-type-specific endosomal compositions and maturation states to develop task-specific delivery strategies. Importantly, such efforts do not necessarily require new CG FF development but rather a more comprehensive application of existing models. Taken together, our findings provide a mechanistic framework for improving the endosomal escape efficiency of RNA-LNP therapeutics and expanding their clinical potential.

## METHODS

### AA-MD Simulations

AA-MD simulations were conducted to generate reference data for SPICA CG modeling of nonbonded interactions between MC3 lipids, dsDNA, and EM components—specifically PS and PI lipids. Simulations were performed using NAMD version 2.12^53^ and GROMACS 2018^54^ with the CHARMM 36 force field^55,56^ and TIP3P water model.^57^ All systems were simulated in the NPT ensemble using a 2 fs integration time step. Electrostatic interactions were computed using the particle-mesh Ewald (PME) method.^58^ Lennard-Jones (LJ) interactions were smoothly truncated using a force-switching function between 10 and 12 Å.

NAMD was specifically used for free energy calculations with the Colvars module.^59,60^ A Langevin thermostat (damping coefficient: 5 ps^-1^) controlled the system temperature at 310 K. Pressure was maintained at 1 atm using the Langevin piston algorithm^61^ with a piston period of 200 and a decay time of 100 fs. Semi-isotropic pressure coupling was applied for simulations of lipid bilayer systems. In GROMACS simulations, temperature and pressure were maintained at 310 K and 1 atm using the Nosé-Hoover thermostat^62,63^ and Parrinello-Rahman barostat,^64^ respectively.

### CG-MD Simulations

CG-MD simulations of LNP, EM, and combined LNP-EM systems were performed using either LAMMPS^65,66^ or GROMACS^54^ with the SPICA force field.^34,37,67^ LJ interactions were truncated at 1.5 nm. Electrostatics were calculated using the particle-particle particle-mesh method in LAMMPS and PME^58^ in GROMACS. System temperature was controlled via the Nosé-Hoover thermostat^62,63^ or a velocity-rescaling method, and pressure was maintained with the Parrinello-Rahman barostat ^64^ using semi-isotropic coupling. A 10 fs integration time step was used.

For the LNP-EM system, simulations included a 1 ns equilibration (NVT ensemble) and a 30 μs production run (N*P*_*xy*_*P*_*z*_T ensemble) at 310 K and 1 atm.

### Surface Tension of Monolayers

To obtain reference data for MC3–PS/PI nonbonded interactions, AA-MD simulations were conducted for binary monolayer systems. MC3 lipids were arranged at a molecular area of 0.80 nm^2^, representing the liquid-phase monolayer. The assumed area per lipid 0.70 nm^2^ for both POPS and POPI. Surface tension convergence in AA-MD was carefully monitored.

Subsequently, CG-MD simulations of equivalent monolayer systems were conducted to reproduce the AA-derived surface tension by optimizing CG nonbonded interaction parameters. The resulting CG surface tension values (listed in Table S1) show good agreement with AA-MD results.

### Density Profile of the CG Segments

AA trajectories of POPS/MC3 and POPI/MC3 systems were mapped to CG representations using SPICA tools (https://github.com/SPICA-group/spica-tools). Density profiles of individual CG segments along the monolayer normal were calculated to further guide parameterization. As shown in Figure S1, the segment distributions from CG-MD closely match those obtained by mapping AA trajectories.

### Free Energy Profile of Nucleobase Permeation

To refine lipid-DNA nonbond interactions, potential of mean force (PMF) profiles were calculated for nucleobases translocation across POPS and POPI bilayers (Fig. S2). For obtaining the PMF, the adaptive biasing force (ABF) method was used, with the reaction coordinate defined along the membrane normal (*z*-axis), where the center of the membrane was set as *z* = 0. The coordinate space was divided into 5 Å windows up to 30 Å from the membrane center. and each window was simulated for at least 300 ns.

### Preparation of the EM-LNP System

A simple EM model composed of 90 POPC, 60 cholesterol, 22 POPE, 14 POPS, and 14 POPI lipids was built using CHARMM-GUI, based on previous reports.^68,69^ This model was converted to CG resolution using SPICA tools and replicated seven times in both x and y dimensions to construct a large planar EM. The LNP contained 6400 MC3 lipids, 4992 cholesterols, 1284 DSPC lipids, and 64 dsDNA molecules (PDB ID: 1BNA), based on prior studies.^34^ In keeping with the acidic environment of the late endosome, all MC3 lipids were modeled in their cationic state. The LNP was placed above the equilibrated EM for subsequent simulations.

### Dehydration of the LNP-Membrane Interface

To initiate fusion, water molecules were removed from the contact region between LNP and EM by applying a spherical wall potential:

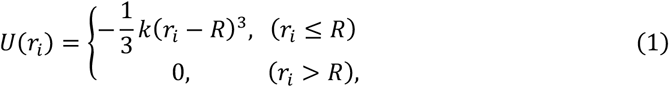

where *r*_*i*_ is the distance of CG water particle *i* from the center of the sphere, *k* is the force constant (10000 kcal/mol/nm^3^), and *R* is the sphere radius (5 nm). The sphere center was positioned at the interface between the LNP and EM. This potential was applied for 100 ns starting from 700 ns into the simulation.

### Angular Distribution of Lipids

To analyze lipid accumulation in the fusion zone (Fig. S6), the tilt angles of EM lipids relative to the membrane normal were computed. Tilt angles were defined as the angle between the lipid head-tail bond vector and the membrane normal, using an in-house Python script. Trajectory data from 1.3 to 1.4 μs were used for this analysis.

### Lateral Tension

To accelerate the membrane fusion, lateral tension was applied to the EM. The surface tension *γ* was computed as:

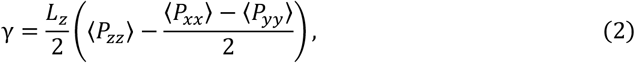

where ⟨*P*_*αα*_⟩ (*α* = *x, y, z*) denotes the ensemble-averaged pressure tensor and *L*_*z*_ is the simulation box length in the *z*-direction. A target surface tension of 4 mN/m was set, consistent with experimental conditions.^70^ At 23.2 μs, *L*_*z*_ was 40.7 nm, requiring *P*_*xx*_ and *P*_*yy*_, to be maintained at −1 bar.

## ACKNOWLEDGEMENTS

This work is supported by JSPS KAKENHI Grant Numbers JP21H01880, JP24H00038, JP24H00843, and JP25K01729. Calculations were performed on the supercomputer facilities of the Institute for Solid State Physics, the University of Tokyo (ISSPkyodo-SC-2023-Ea-0002, 2023-Eb-0004, 2024-Ea-0001, 2024-Eb-0003, 2025-Ea-0002), and the Research Center for Computational Science, Okazaki, Japan (Project: 25-IMS-C094, 24-IMS-C090, 23-IMS-C095).

## Author Contributions

A.P.S. led the data analysis, figure preparation, and interpretation of results, and wrote the main draft of the manuscript. K.S. performed the molecular dynamics simulations and initial data analysis. Y.M. contributed to the revision and improvement of the figures. W.S. supervised the project, provided critical input on data interpretation, and substantially revised the manuscript. All authors discussed the results and contributed to the final version of the manuscript.

## Data Availability

The data collected and analyzed in this research paper are available on Zenodo (https://zenodo.org/records/17489583). Additionally, more information about the SPICA force field, including tutorials and parameter files, can be found at http://www.spica-ff.org.

## Competing Interests

The authors declare no competing interests.

## Notes

### Competing Interest Statement

The authors have declared no competing interest.

