## Supplementary Information for "pH-Sensitive Amino Lipid-Driven Pore Formation Enables Endosomal Escape of Lipid Nanoparticles"

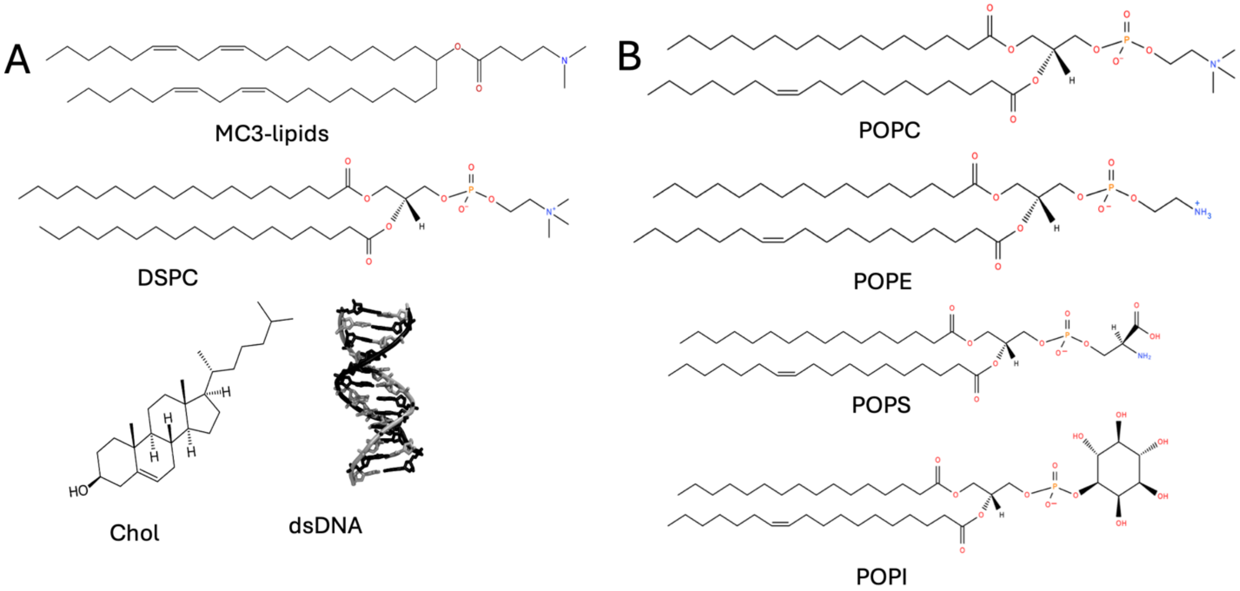


Figure S1. (*A, B*) Molecular structures of the LNP and EM components.

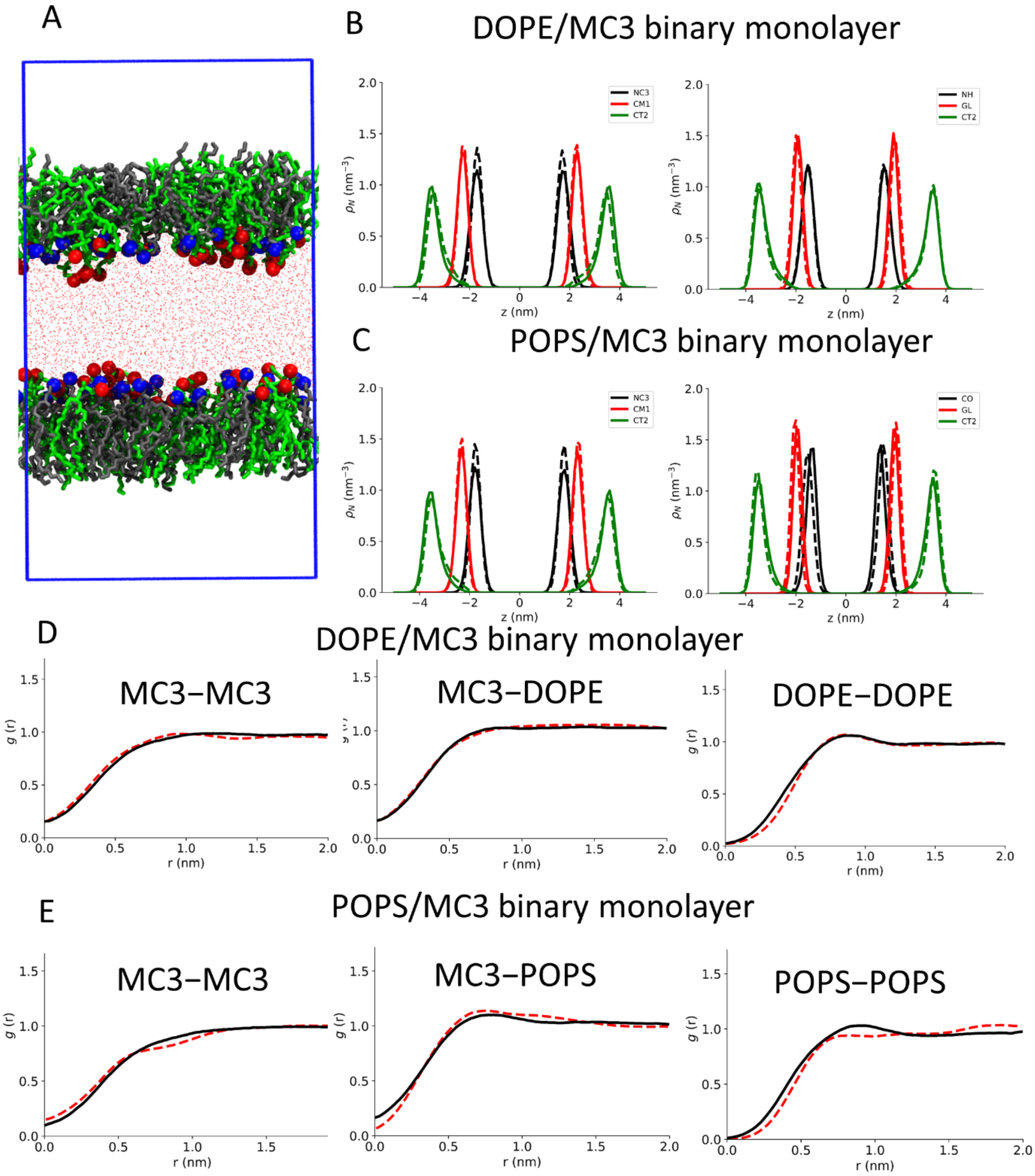


**Figure S2.** *(A)* Snapshot of the all-atom (AA) simulation box for the cationic MC3 lipid/DOPE monolayer. Color scheme: grey for MC3, green for DOPE, blue spheres for head-group sites of MC3, red spheres for head-group sites of DOPE, and red dots represent water oxygen. *(B, C)* Number density profiles of selected CG segments along the monolayer normal of the *(B)* cationic MC3/DOPE and *(C)* MC3/POPS binary systems. Solid and dashed lines represent CG- and AA-MD results, respectively. *(D, E)* Two-dimensional radial distribution functions (2D-RDFs) between lipids in the *(D)* MC3/DOPE and *(E)* MC3/POPS binary monolayers, confirming the accuracy of the CG parameters. The red dotted and black solid lines represent the 2D-RDFs obtained from AA- and CG-MD, respectively.

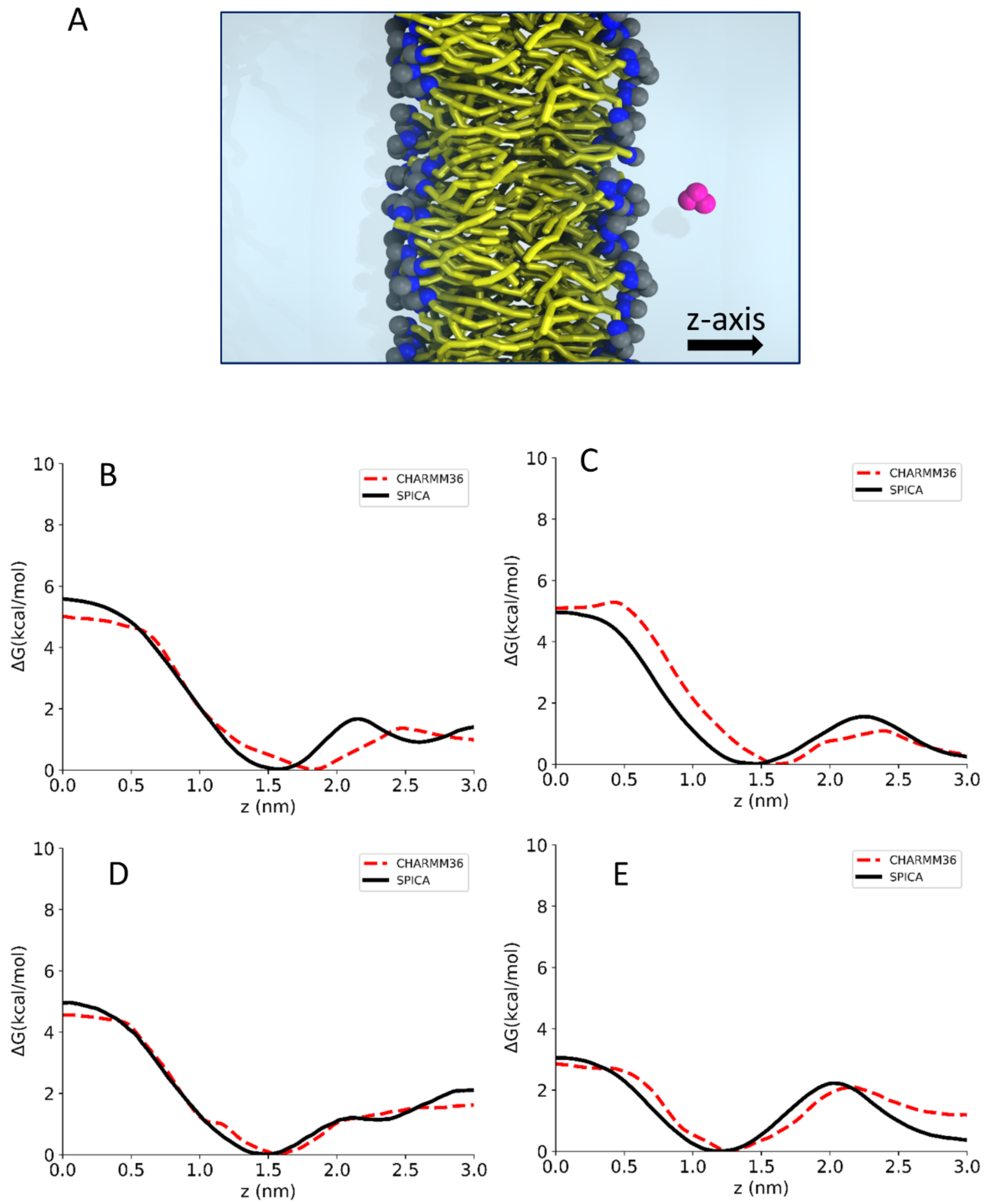


**Figure S3.** *(A)* Representative snapshot of the CG-MD simulation performed to obtain the partition-free energy of nucleic acid analogs from an POPS lipid monolayer. Color scheme: yellow for POPC tails, blue and gray spheres for head-group sites of POPS, ice blue for water phase and magenta for thymine. *(B, D)* Free energy profiles of thymine as a function of distance from the center of the *(B)* POPS and *(D)* POPI bilayer along the bilayer normal (*z*-axis). *(C, E)* Free energy profiles of deoxyribose as a function of distance from the center of the *(C)* POPS and *(E)* POPI bilayer along the bilayer normal (*z*-axis). Color scheme: red dashed lines represent AA-MD results using the CHARMM36 force field (FF), and black solid lines represent CG-MD results using the optimized SPICA FF.

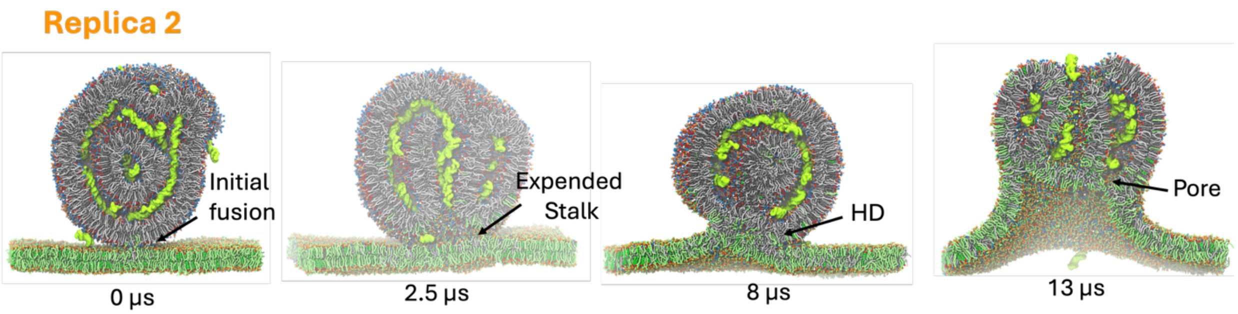


**Figure S4. Representative snapshots of *Replica 2* illustrate the time evolution of the LNP-EM system during molecular dynamics (MD) simulations.** These snapshots demonstrate that the radial expansion of the stalk and the subsequent formation of the hemifusion diaphragm (HD) lead to the initiation of a small fusion pore, similar to what is observed in *Replica 1*. Color scheme: Different lipid types are shown in distinct colors. Lipid headgroups, tails, and dsDNA molecules are depicted using van der Waals (VDW), Licorice, and QuickSurf (an iso-surface) representation, respectively. For clarity, water and ions are omitted.

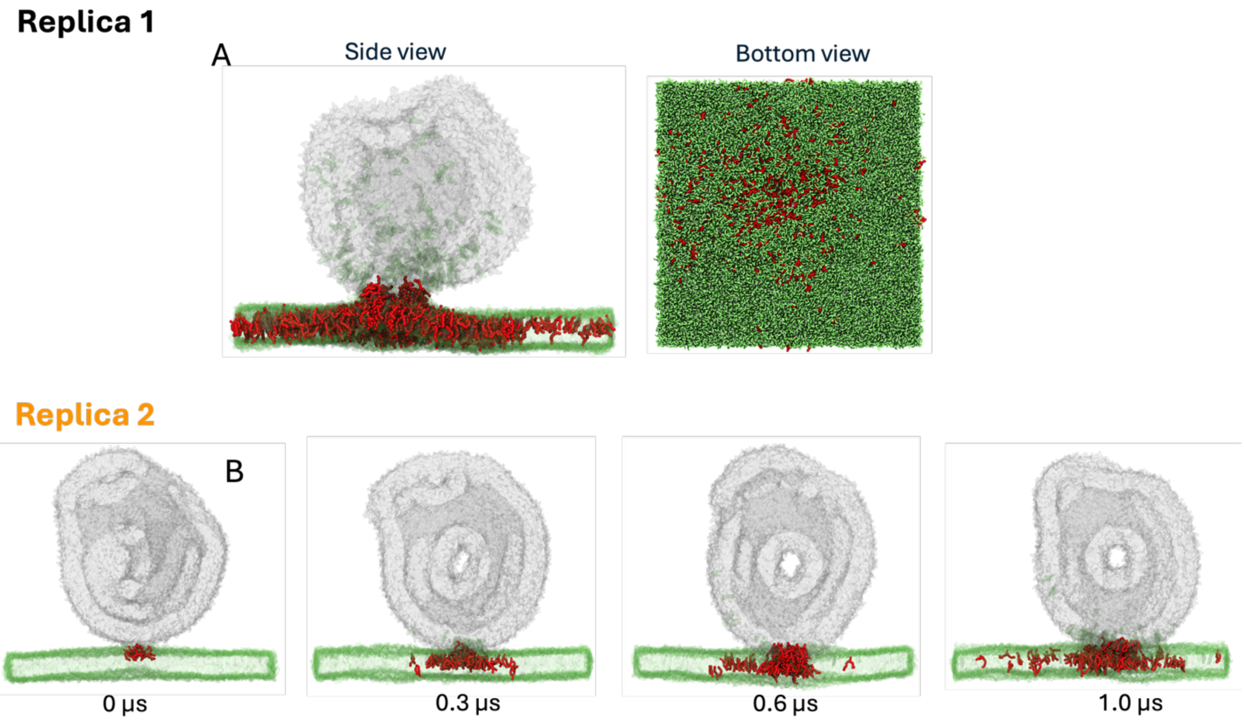


**Figure S5. MC3-lipids exhibit rapid migration in the endosomal membrane (EM)**. *(A)* The cross-sectional view and bottom view of the representative MD simulation snapshot of the LNP-EM complex (*Replica 1*) at t ~ 0.9 μs. Color scheme: red for MC3-lipids, gray and green represent the LNP and EM domains, respectively. **(*B*)** The time evaluation of the LNP-EM complex simulation in *Replica 2* also highlighted MC3-lipids migration in EM. The color scheme is identical to that of Figure S4B.

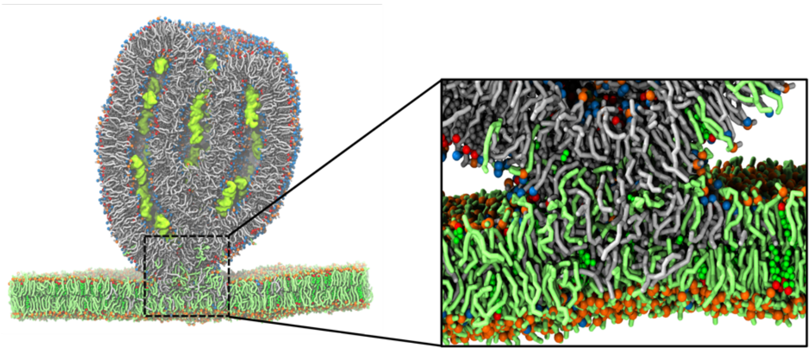

**Figure S6. A disrupted stalk compared to conventional lipid bilayer/vesicles and membrane fusion stalk.**[1] The cross-sectional view of representative MD simulation snapshot of the LNP-EM complex after fusion at t ~ 0.25 μs. Color scheme: Different lipid types are shown in distinct colors. Lipid headgroups, tails, and dsDNA molecules are depicted using van der Waals (VDW), Licorice, and QuickSurf (an iso-surface) representation, respectively. For clarity, water and ions are omitted. Inset: zoom of the fusion region for comparison with previously reported bilayer/vesicle and membrane fusion.[1]

[1]Kawamoto S., Klein, M. L., & Shinoda W. Coarse-grained molecular dynamics study of membrane fusion: Curvature effects on free energy barriers along the stalk mechanism, *J. Chem. Phys.* **143**, 243112 (2015)

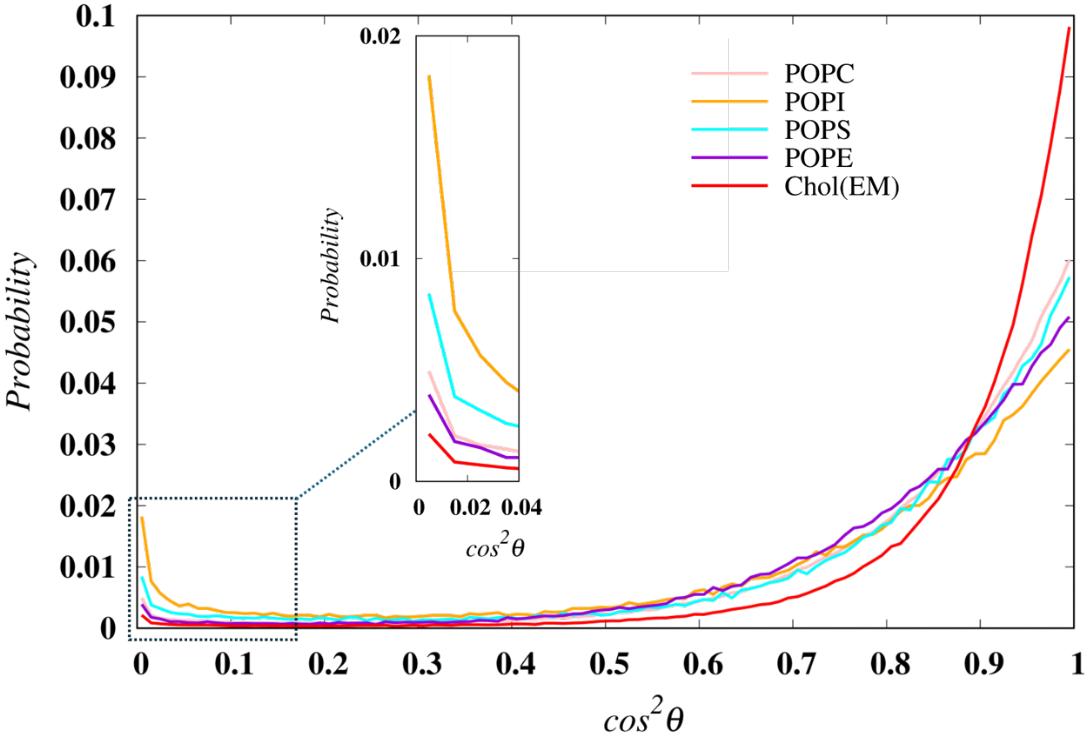


**Figure S7. The orientation of lipids in relation to the membrane's normal direction emphasizes their tendency to migrate toward LNPs**. Probability distribution of the angle, $\cos^2\theta$, between the lipid head-to-tail bond vector and the membrane normal. Inset: Distribution of $\cos^2\theta$ in the range $0 \leq \cos^2\theta \leq 0.04$.

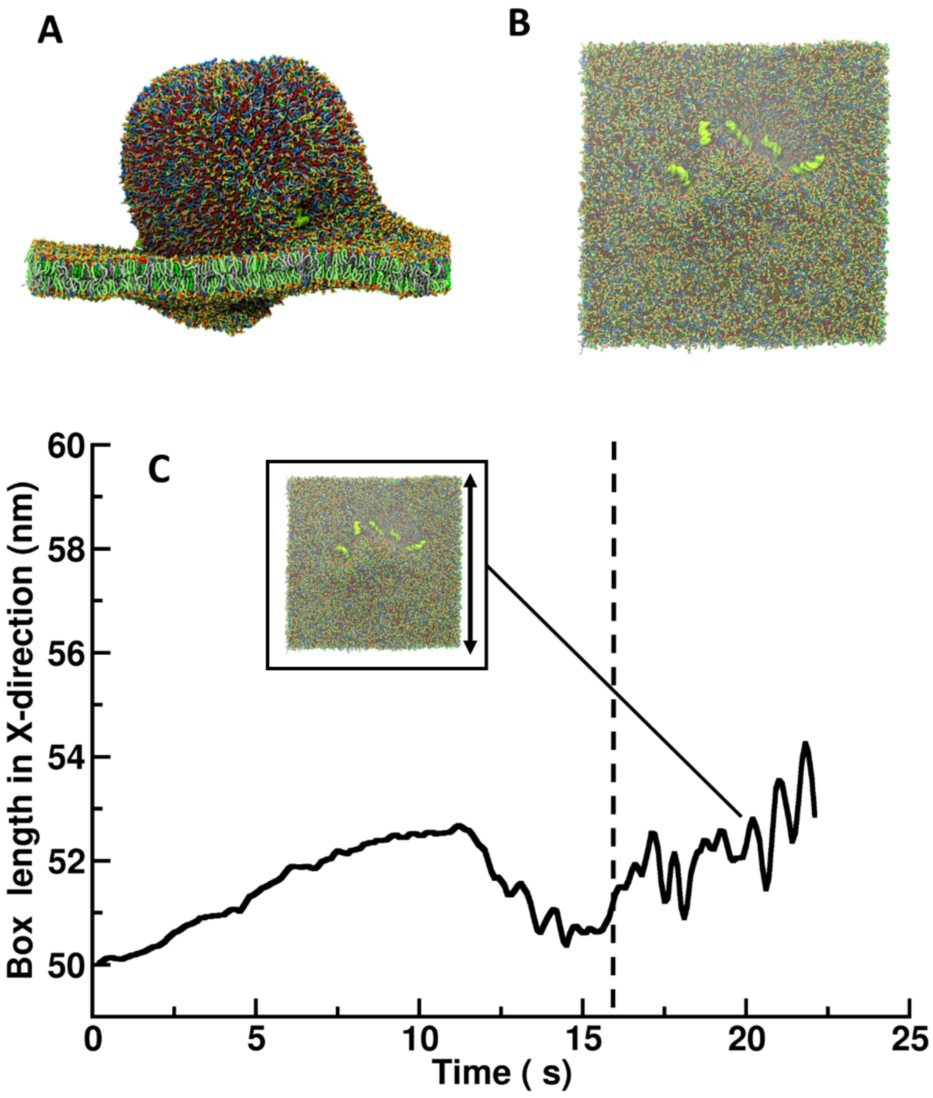


**Figure S8.** Representative snapshots of the EM-LNP fused complex containing all MC3 lipids in protonated state at 18.4 μs: *(A)* side view and *(B)* top view. *(C)* Time evolution of the box length in the X-direction during the MD simulations.

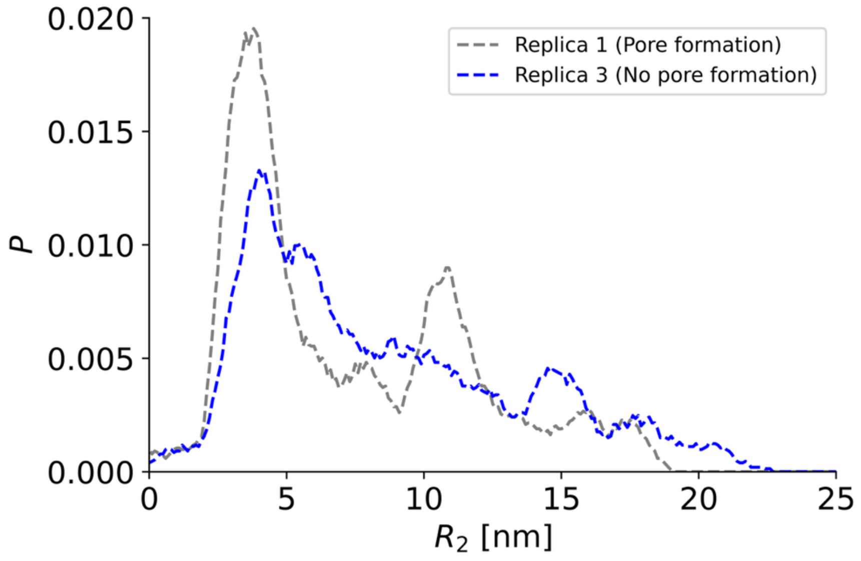


**Figure S9** Probability distributions of dsDNA in the LNP-EM fused complex at ~8 μs, along the EM surface normal from the EM center toward the LNP interior. The distance from the EM center, $R_2$, is defined as the inward projection of $R_2 = R_i - 2r_{SASA}$ onto the EM surface normal, shifted by twice the radius of the SASA probe particle.

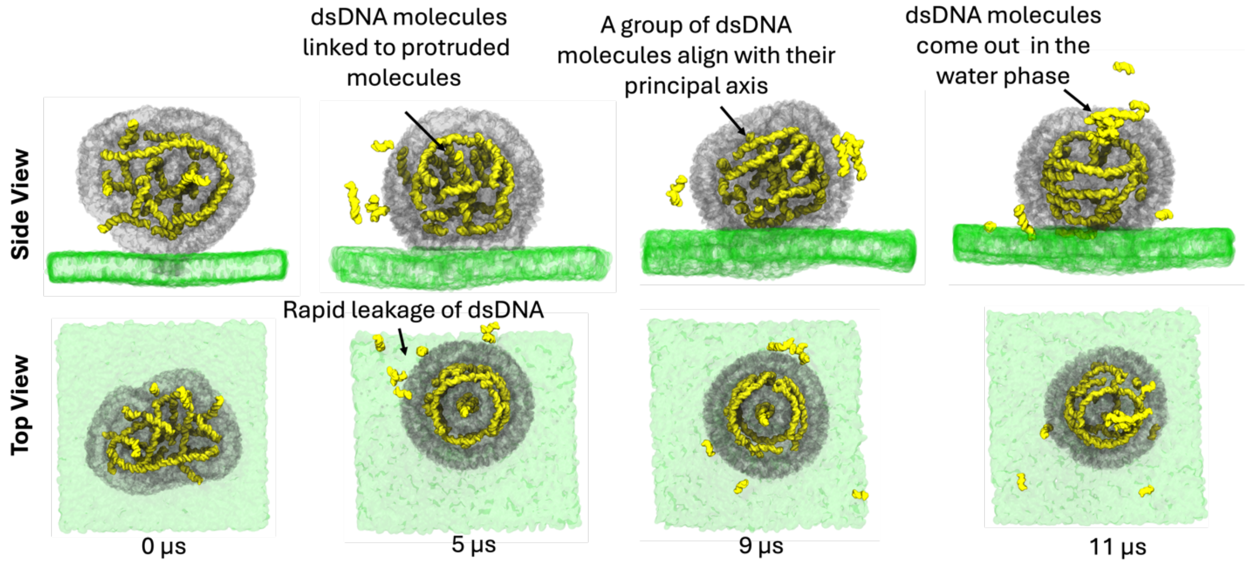


**Figure S10.** illustrates the time evolution of the system without pore formation *(Replica 3)*, highlighting two main observations: *(i)* the rapid leakage of double-stranded DNA (dsDNA), and *(ii)* dsDNA molecules linked to protruding molecules from the lipid nanoparticle (LNP) surface. Subsequently, these dsDNA molecules align with the principal axis of the LNP and migrate into the water phase of luminal side. Color scheme: yellow represents dsDNA molecules, gray and green indicate the LNP and EM domains, respectively. The results of *Replica 4* have been omitted as they are similar to those presented here.

**Table S1**. Comparison of surface tension ($\gamma$, in mN/m) for binary monolayer systems between AA- and CG-MD simulations.

| Monolayer system* | $\gamma$ (AA MD) | $\gamma$ (CG MD) |
|---|---|---|
| cationic MC3/DOPE (1:1) | 46 ± 1 | 44 ± 1 |
| cationic MC3/POPS (1:1) | 52 ± 1 | 50 ± 1 |
| cationic MC3/POPI (1:1) | 58 ± 1 | 55 ± 1 |

**Table S2.** Molecular composition of the LNP-EM complex systems simulated in this study.

| | | | Molecule | Number |
|---|---|---|---|---|
| LNP | pH ~ 4 | Replica 1[#] | cationic-MC3 | 6,400 |
| | | | Chol | 4,992 |
| | | | DSPC | 1,284 |
| | | | dsDNA | 64 |
| | pH ~ 5 | Replica 2[#]<br>Replica 3*<br>Replica 4* | cationic-MC3 | 5,174 |
| | | | neutral-MC3 | 1,226 |
| | | | Chol | 4,992 |
| | | | DSPC | 1,284 |
| | | | dsDNA | 64 |
| | | EM | POPC | 4,410 |
| | | | Chol | 2,940 |
| | | | POPE | 1,078 |
| | | | POPI | 686 |
| | | | POPS | 686 |
| | | Solvent | WAT | ~1,350,000 |

[#]*Replica* 1 and 2 that included endosomal membrane (EM) exhibited pore formation. In contrast, **Replica* 3 and 4 with EM did not show any pore formation during a runtime of 20 to 30 μs. All systems feature explicit coarse-grained (CG) water and sodium/chloride (NaCl) counter-ions. Additionally, *Replica* 2, 3, and 4 contain physiological ion concentrations of 150 mM NaCl.

**Table S3.** Number of lipids exchanged between the LNP and the EM at 1.4 μs

| System | Lipids | Total number of molecules | LNP (remained) (moved) | EM (moved) (remained) | Exchanged lipid ratio (%) |
|---|---|---|---|---|---|
| LNP | MC3 | 6400 | 5574 | 826 | 12.9 |
| | Chol (LNP) | 4992 | 4642 | 350 | 7.0 |
| | DSPC | 1284 | 1247 | 37 | 2.9 |
| EM | POPI | 686 | 89 | 597 | 13.0 |
| | POPS | 686 | 78 | 608 | 11.4 |
| | POPC | 4410 | 240 | 4170 | 5.4 |
| | POPE | 1078 | 54 | 1024 | 5.0 |
| | Chol (EM) | 2940 | 61 | 2879 | 2.1 |
| Total | | | 11985 | 10491 | |
